# Tissue fibroblasts are a critical source of prostacyclin and anti-thrombotic protection

**DOI:** 10.1101/2022.01.11.475814

**Authors:** Jane A. Mitchell, Maria Vinokurova, Maria Elisa Lopes-Pires, Fisnik Shala, Paul C. Armstrong, Blerina Ahmetaj-Shala, Youssef Elghazouli, Bin Liu, Yingbi Zhou, Chuan-ming Hao, Harvey R. Herschman, Nicholas S. Kirkby

## Abstract

Prostacyclin is one of the bodies fundamental signalling pathways. It has long been considered principally anti-thrombotic hormone derived from the vascular endothelium. The role of non-vascular sources in prostacyclin synthesis has not been systematically evaluated, however, due to a lack of tools resulting in an underappreciation of its role in other contexts. Here we used cellspecific knockout mice and human tissues to show that lung, and other tissues, are powerful producers of prostacyclin independent of their vascular components. Instead, in mice and humans, lung prostacyclin synthesis is associated with fibroblasts. The fibroblast-derived prostaglandins enter the circulation and provide systemic anti-thrombotic protection. These observations define a new paradigm in prostacyclin biology in which fibroblast/non-vascular-derived prostacyclin works in parallel with endothelium-derived prostaglandins to control cardiovascular health and potentially a broad range of other biology. These results may explain how local diseases of the lung and elsewhere result in cardiovascular risk.

## Introduction

Prostacyclin is the most potent and powerful endogenous inhibitor of platelet activation and represents one of the bodies fundamental anti-thrombotic and cardioprotective pathways(Mitchell and Kirkby, 2019; Moncada et al., 1976). This anti-platelet activity is principally mediated by activation of the IP receptor. IP is highly expressed on platelets; IP activation elicits elevation of intra-platelet cyclic AMP levels. Consequently, IP inhibition or genetic deficiency results in loss of sensitivity to exogenous prostacyclin, a pro-thrombotic phenotype in animal models(Mitchell et al., 2019; Murata et al., 1997) and an increased risk of heart attacks and strokes in man(Arehart et al., 2008). Prostacyclin can also, in some contexts, act as a vasodilator(Moncada *et al*., 1976) and protect against atherogenesis(Kobayashi et al., 2004). Dysfunction of the prostacyclin pathway is a feature of several vascular pathologies, including pulmonary hypertension where exogenous prostacyclin is an established therapy(Mitchell and Kirkby, 2019). In addition to its cardiovascular actions, prostacyclin is important in lung, gastrointestinal and renal function, in pain/inflammation, and in the regulation of the immune system. As such, understanding prostacyclin biology is essential not only for cardiovascular health, but also for the proper functioning of a broad range of organ systems.

Prostacyclin is produced within a larger family of prostanoid mediators by the activities of phospholipases, cyclo-oxygenases and prostacyclin synthase, through step-wise metabolism of membrane lipids (Mitchell and Kirkby, 2019). Phospholipases liberate the substrate, arachidonic acid, from cell membranes. Although at least 18 other isoforms of phospholipase A2 exist, arachidonic acid for prostacyclin synthesis is principally released from membranes by cytosolic phospholipase A2(Kirkby et al., 2015). However, other phospholipases, as well as members of the phospholipase C family, have also been implicated in prostacyclin production. Phospholipase enzymes respond to elevated Ca^2+^ or other acute signals, conferring a requirement for specific stimulation for prostanoid production. Cyclo-oxygenases, which exist as two isoforms, convert the phospholipase-derived arachidonic acid to an unstable intermediate, prostaglandin H_2_. Cyclo-oxygenase-1 is, like cytosolic phospholipase A2, widely expressed as a physiological house-keeping enzyme; in contrast, constitutive cyclo-oxyenase-2 expression is restricted to certain regions that include cells of the kidney, brain and gut(Kirkby et al., 2013b). However, cyclo-oxyenase-2 can be rapidly induced in many other tissues during inflammation and proliferation. The relative importance of cyclooxygenase-1 and −2 in prostacyclin production has remained a controversial area. Cyclo-oxygenase-2 is associated with prostacyclin metabolites in the urine(McAdam et al., 1999); however, this may reflect renal rather than systemic production(Mitchell et al., 2018). Cyclo-oxyenase-2 also plays a role in prostacyclin production during gross systemic inflammation(Kirkby et al., 2013a). Where studied directly, however, most evidence supports cyclo-oxygenase-1 as the principal cyclooxygenase isoform driving bulk constitutive prostacyclin production in mouse (Kirkby et al., 2012; Liu et al., 2012b) and human cells/tissues(Mitchell et al., 2006). Notably, mice deficient in cyclo-oxygenase-1, but not cyclo-oxygenase-2, show a profound loss of prostacyclin generation across almost all tissues in the body(Kirkby *et al*., 2012; Liu *et al*., 2012b).

The final step in prostacyclin synthesis is conversion of prostaglandin H_2_ to prostacyclin by the enzyme prostacyclin synthase. In contrast to the other enzymes in the pathway, prostacyclin synthase has a narrower cell and tissue expression pattern, resulting in spatial differences in prostacyclin production by different cells and tissues. Prostacyclin was first discovered as a substance released from the endothelial cell layer of arterial tissue (Mitchell *et al*., 2019; Moncada et al., 1977). This observation has been reproduced countless times; arterial tissue ex vivo and arterial endothelial cells in vitro have been universally observed to possess a robust capacity for prostacyclin synthesis. By comparison, platelets(Moncada *et al*., 1976) and leucocytes(Kennedy et al., 1980) are almost entirely deficient in their ability to produce prostacyclin under physiological conditions. After its discovery, it was quickly realised that prostacyclin was also produced by isolated tissues; it is abundant in tissue perfusates and was found to be the major prostanoid generated by the lung and the heart(de Deckere et al., 1977; Gryglewski et al., 1978). Although in vitro evidence has indicated many cell types, including epithelial cells(Taylor et al., 1979), fibroblasts(Claesson et al., 1977) and smooth muscle cells (Baenziger et al., 1979), have at least some prostacyclin synthetic capacity, the consensus in the field has remained that, within tissues, the bulk of prostacyclin production is associated with their vascular compartments. The validity of this assumption has been difficult to test, because when individual cells are cultured they undergo rapid alterations in prostanoid synthetic pathways(Ager et al., 1982). Moreover, there have been no tools available that allow the contribution of individual cell types to be assessed for prostacyclin production in intact tissues or in vivo. We have recently described mice in which cyclo-oxygeanse-1, cyclo-oxygenase −2(Mitchell *et al*., 2019), or prostacyclin synthase(Cao et al., 2019) can be deleted specifically from vascular endothelial cells. These mouse models provide new tools to foster the understanding of the cellular origins of prostacyclin within tissues. In the current study we use them to determine whether prostacyclin synthesis is purely a product of the vasculature, or whether there are additional, relatively neglected depots of prostacyclin production in the body that contribute to local organ function and/or systemic anti-thrombotic protection.

## Results

### Tissues are major sources of prostacyclin generated through cyclo-oxygenase-1

The view that prostacyclin is a principally a product of blood vessels has become a firmly fixed tenant of cardiovascular biology. Whilst all tissues are capable of producing prostacyclin, the relative prostacyclin generating capacity of different tissues is not fully appreciated. Using the aorta as a benchmark, we assayed prostacyclin formation per unit mass from paired mouse tissues stimulated with Ca^2+^ ionophore to maximally activate endogenous prostacyclin production. Prostacyclin release (measured as 6-keto-PGF_1α_) was observed from all tissues, with the lowest release from the renal cortex and heart and the highest release from the lung and colon (Figure 1A). Lung and colon produced equivalent prostacyclin to aorta on a ‘per mg tissue’ basis. Considering the large total mass of these tissues suggests that they (and their constituent vascular components) may be major contributors to whole body prostacyclin generation. The relative ability of tissues and their included vessels to generate prostacyclin broadly correlated with prostacyclin synthase gene (*Ptgis*) expression; expression of this enzyme is enriched in aorta, colon and lung relative to other tissues (Figure 1B). By contrast, relative tissue levels of cyclo-oxygenase-1 gene (*Ptgs1*) expression correlated poorly with prostacyclin release; for example, the aorta and lung expressing comparatively low levels of the *Ptgs1* gene (Figure 1B). Cyclo-oxygeanse-2 (*Ptgs2*) was very weakly expressed across all tissues (~10-100-fold less than *Ptgs1*; Figure 1B) in keeping with our previous observations of the relative constitutive expression and activity of the two cyclo-oxygenase isoforms(Kirkby *et al*., 2012). In agreement, global cyclo-oxygenase-1 (*Ptgs1*^-/-^) deletion abolished prostacyclin generation both in aortae and in all tissues studied (Figure 1C). Thus, arteries and tissues both generate prostacyclin which is (i) driven by cyclo-oxygenase-1 activity but (ii) at a gene expression level, reflects the relative level of prostaglandin synthase.

**Figure 1.**
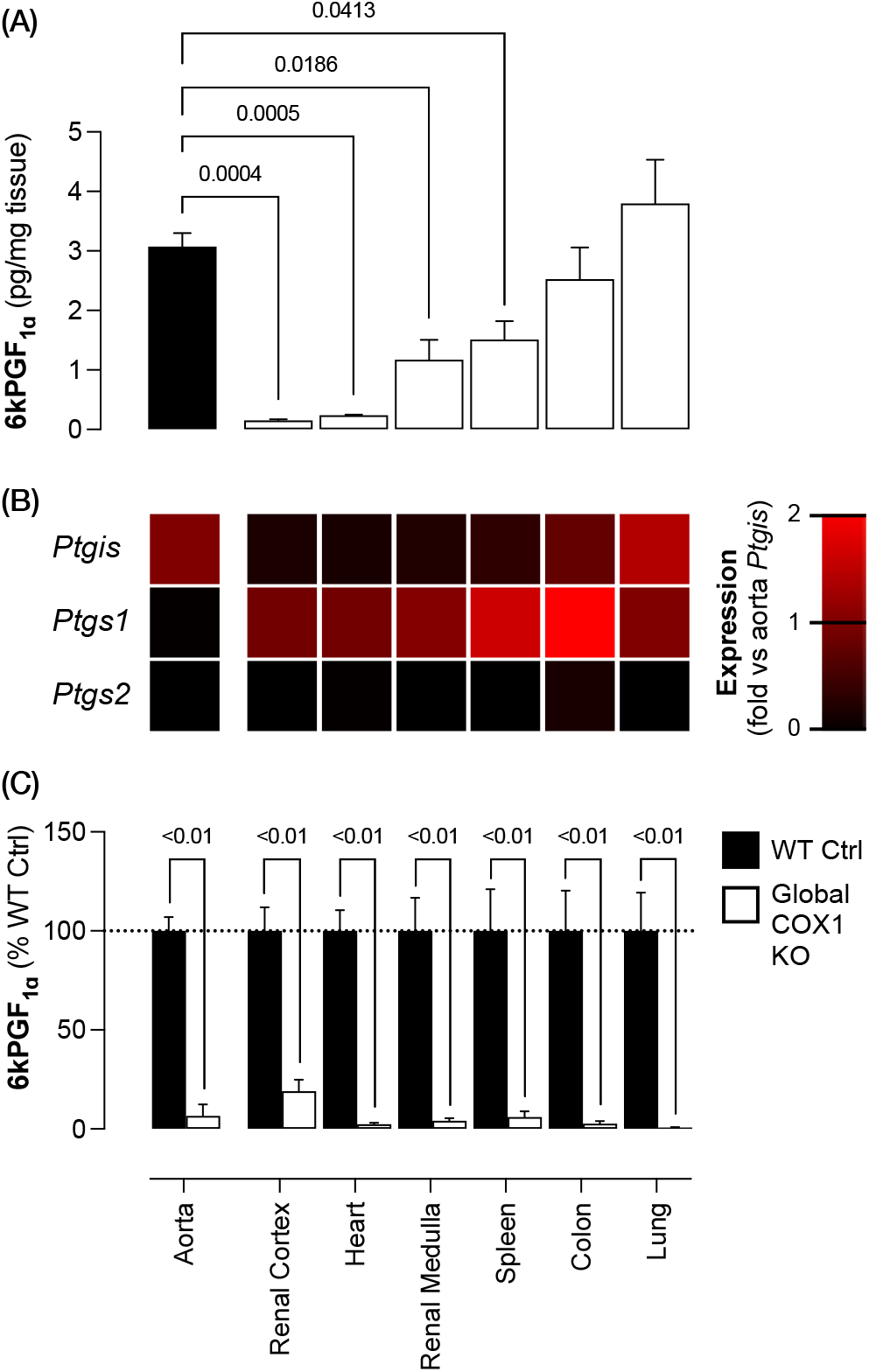
Both tissues and arteries generate prostacyclin which reflects relative prostacyclin synthase expression and requires cyclo-oxygenase-1. (A) Prostacyclin levels (measured as 6kPGFiα after A23187 Ca^2+^ ionophore 30μM stimulation) per unit mass (n=4) and (B) *Ptgis* (prostacyclin synthase), *Ptgs1* (cyclo-oxygenase-1) and *Ptgs2* (cyclo-oxygenase-2) gene expression (n=4) in wild-type mouse aorta and tissue. (C) Prostacyclin release from tissues from global cyclo-oxygenase-1 knockout (global COX-1 KO) and matched wild-type control (WT Ctrl) mice (n=3-6). Data are mean ± SEM with p values by one-way ANOVA (A) or unpaired t-test (C) indicated where p<0.05.

### Many tissues can produce prostacyclin in the absence of endothelial cell cyclo-oxygenase-1

To understand the nature of tissue prostacyclin generation, we next considered whether the ability to release prostacyclin is simply a function of the tissues constituent endothelial component, or whether non-vascular cell types within complex tissues generate prostacyclin directly. To do this we used mice in which cyclo-oxygenase-1 is specifically deleted from endothelial cells (Ptgs1^flox/flox^; VE-cadherin-Cre^ERT2^). These mice have been characterised previously(Mitchell *et al*., 2019). As we have seen before, aortic rings from these mice had marked reduction (~80%) in prostacyclin release (Figure 2A). Here we extended this observation to show a consistent loss of prostacyclin synthesis in veins (Figure 2B) and arteries supplying the lung (Figure 2C), kidney (Figure 2D) and gut (Figure 2E) from endothelial cyclo-oxygenase-1-deficient mice. However, the effect of endothelial cyclo-oxygenase-1 deletion on prostacyclin release from other isolated tissue segments was variable. In the heart, prostacyclin release when endothelial cyclo-oxygenase 1 is deleted was reduced ~50% suggesting a major role of endothelial cyclo-oxygnease-1 in prostacyclin in this tissue (Figure 2F). In contrast, in the lung (Figure 2G), colon (Figure 2H), kidney (Figure 2I) and spleen (Figure 2J) endothelial cyclo-oxygenase-1 deletion had no effect on prostacyclin release. The residual ‘endothelium-independent’ tissue prostacyclin release was not accounted for by vascular smooth muscle cyclo-oxygenase-1 activity; smooth muscle cyclo-oxygenase-1 knockout mice (*Ptgs1*^flox/flox^;Smmhc-Cre), which we have previously characterised(Mitchell *et al*., 2019), exhibited no change in prostacyclin release in any tissue studied (Figure 2F-J). Thus, whilst in large arteries and veins prostacyclin production is almost entirely dependent on prostaglandin H_2_ generated by vascular endothelial cyclo-oxygenase-1, within most tissues, prostacyclin production appears to be driven by non-vascular cyclo-oxygenase-1 activity present elsewhere.

**Figure 2.**
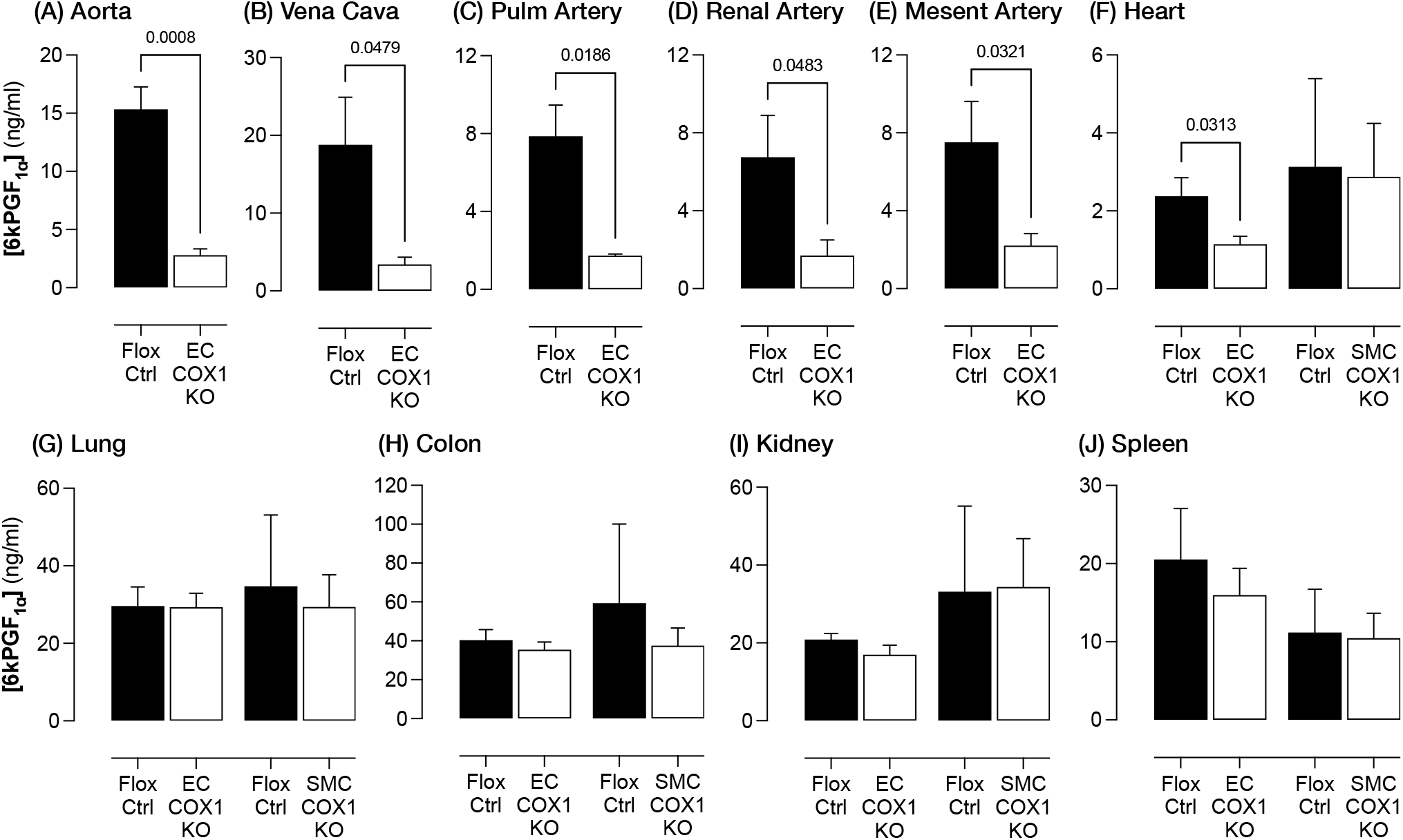
Role of vascular cyclo-oxygenase-1 in prostacyclin release from arteries, veins and tissues. Prostacyclin release (measured as 6kPGF_1α_ after A23187 Ca^2+^ ionophore 30 μM stimulation) from (A) isolated aorta (n=4), (B) vena cava (n=4), (C) pulmonary artery (n=3), (D) renal artery (n=7) and (E) mesenteric artery (n=7) from endothelial cyclo-oxygenase-1 knockout (EC COX1 KO) and floxed littermate control animals (Flox Ctrl), and from intact segments of (F) heart (left ventricle; n=5-8), (G) lung (parenchyma; n=5-8), (H) colon (n=5-8), (I) kidney (renal medulla; n=5-8) and (J) spleen (n=5-8) from EC COX1 KO mice, smooth muscle cyclo-oxygenase-1 knockout (SMC COX1 KO) and their respective floxed littermate controls (Flox Ctrl). Data are mean ± SEM with p values by unpaired t-test indicated where p<0.05.

To validate and explore further these observations we focused on the lung because (1) lung has the highest prostaglandin production amongst tissues (Figure 1A), (2) lung prostacyclin production appeared to be almost completely *independent* of vascular cyclo-oxygenase-1 activity (Figure 2G) and (3) prostacyclin generated in the lung has been suggested to act directly on the heart and arterial vessels due to their anatomical arrangement(Gryglewski *et al*., 1978). With no apparent requirement for endothelial/vascular smooth muscle cyclo-oxygenase activity-1 in lung prostacyclin production, we first considered if there may be other sources that can donate cyclo-oxygenase-1 generated prostaglandin H_2_ to endothelial cells for conversion to mature prostacyclin, bypassing the effect of endothelial cyclo-oxygenase deletion. This phenomenon of ‘transcellular metabolism’ has been previously described in platelet/endothelial co-cultures where platelet-derived prostaglandin H_2_ can enter endothelial cells to access prostacyclin synthase(Marcus et al., 1980). To address this possibility, we repeated lung prostacyclin release experiments with endothelial cyclo-oxygenase-1 knockout mouse tissue after flushing the pulmonary vasculature of blood to remove circulating platelets. In lung tissue cleared of blood, endothelial cyclo-oxygenase-1 deletion sill had no effect on prostacyclin release (Figure 3A). We further confirmed this result by studying a dual endothelial/platelet cyclo-oxygenase-1 knockout mouse (*Ptgs1*^flox/flox^;Tie2-Cre) where prostaglandin H_2_ cannot be synthesised either by platelets or by endothelial cells(Mitchell *et al*., 2019). As observed in endothelial-specific cyclo-oxygenase-1 knockout mice, endothelial/platelet cyclo-oxygenase-1 deletion had no effect on lung tissue release of prostacyclin stimulated by Ca^2+^ ionophore (Figure 3B). Similarly, when endogenous phospholipase A2 was bypassed with exogenous arachidonic acid there was no effect on prostacyclin production (Figure 3C). To confirm that this observation was not an artifact associated with ex vivo tissue stimulation we also measured prostacyclin levels in unstimulated snap frozen lung tissue homogenates, to estimate levels in the lung *in vivo*. We again found no effect of endothelial/platelet cyclo-oxygenase-1 deletion (Figure 3D). Endothelial/platelet cyclo-oxygenase-1 deletion also had no effect on stimulated release of prostaglandin E2, the other major prostanoid product of the lung (Flox Ctrl: 2.0±0.6ng/ml; EC/PT COX1 KO: 1.9±0.7; n=4; p>0.05 by unpaired t-test). To firmly exclude a role for other pathways of prostaglandin H_2_ generation, we studied lung from endothelial/platelet prostacyclin synthase knockout mice (Figure 3E; *Ptgis*^flox/flox^;Tie2-Cre) and endothelial/platelet cyclo-oxygenase-2 knockout mice (Figure 3F; *Ptgs2*^flox/flox^;Tie2-Cre). As observed for endothelial cyclo-oxygenase-1-deficient models, lung from both strains retained a full capacity to generate prostacyclin; the results confirm that endothelial cells in the lung are neither required to generate prostaglandin H_2_ to support prostacyclin synthesis, nor to convert prostaglandin H_2_ from other sources into prostacyclin. Additional data from these mouse knockout models also corroborated the relative role of endothelium in prostacyclin release in other tissues – these are presented in Supplementary Table 1 but not discussed further.

**Figure 3.**
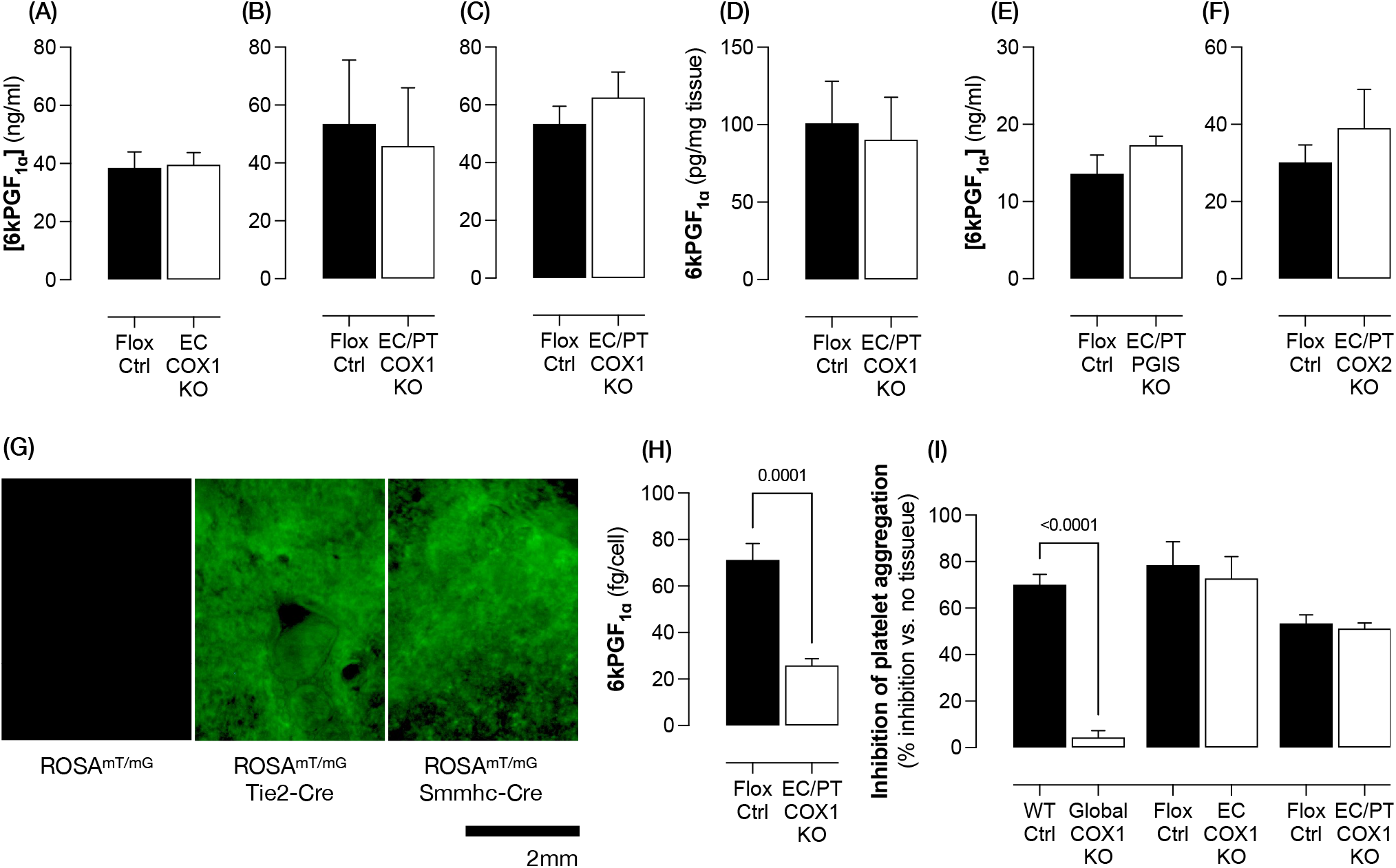
Lung prostacyclin production does not require cyclo-oyxgenase-1, cyclo-oxgenase-2 or prostacyclin synthase in endothelial cells or platelets. (A) Prostacyclin release (measured as 6kPGF_1α_ after A23187 Ca^2+^ ionophore 30μM stimulation) from lung parenchyma segments from endothelial cyclo-oxygenase-1 knockout (EC COX1 KO) and floxed littermate control animals (Flox Ctrl) in which the lung vasculature has been flushed of blood (n=5-6). (B) Prostacyclin from lung parenchyma segments stimulated after stimulation with A23187 Ca^2+^ ionophore (n=6; 30μM) or (C) arachidonic acid (n=6; 30μM) or (D) levels in unstimulated snap frozen lung homogenates (n=6) from endothelial/platelet cyclo-oxygenase-1 knockout mice (EC/PT COX1 KO). Prostacyclin release (A23187 Ca^2+^ ionophore 30μM stimulation) from lung parenchyma segments from (E) endothelial/platelet prostacyclin synthase knockout mice (EC/PT PGIS KO; n=7-13) and (F) endothelial/platelet cyclo-oxygnease-2 knockout mice (EC/PT COX2 KO; n=5), each compared to respective floxed littermate controls. (G) EGFP (green) fluorescence in lung segments of ROSA^mT/mG^ mice with/without a Tie2-Cre or Smmhc-Cre transgene (representative of n=3/genotype). (H) Prostacyclin release from endothelial cells (CD31^+^, CD45^-^, CD41^-^) isolated by FACS from EC/PT COX1 KO and Flox Ctrl mouse lung (n=6). (I) Bioassay of prostacyclin activity as inhibition of human platelet aggregation by lung parenchyma segments from global cyclo-oxygeanse-1 knockout mice (Global COX1 KO) and matched wild-type controls (WT Ctrl) or from endothelial cyclo-oxygenase-1 knockout mice (EC COX1 KO), EC/PT COX1 KO and their respective floxed littermate controls (Flox Ctrl) (n=3-5). Data are mean ± SEM with p values by unpaired t-test indicated where p<0.05.

We next considered whether retention of prostacyclin generation by the lung in these models might be due to incomplete penetrance of Cre-mediated recombination in tissue versus arterial endothelial cells. Previous studies have effectively used Tie2-Cre(Boettcher et al., 2014) and VE-cadherin-Cre(Konishi et al., 2017) to delete floxed genes in lung microvasculature. To confirm this phenomenon in our own animals we crossed Tie2-Cre mice (used to generated endothelial/platelet cyclo-oxygenase-1, cyclo-oxygenase-2 and prostacyclin synthase knockout mice) and Smmhc-Cre mice (used to generate smooth muscle cyclo-oxygenase-1 knockout mice) with an EGFP reporter strain. Robust recombination occurred throughout the lung (Figure 3G). We went on to isolate live microvascular endothelial cells from lung of endothelial/platelet cyclo-oxygenase-1 knockout mice by FACS and found ~70% reduction in prostacyclin production compared to cells isolated from the lung of floxed littermate control mice; these data confirm effective loss of cyclo-oxygenase-1 activity in lung endothelium (Figure 3H). Finally, to ensure that 6keto-PGF_1α_ detected by immunoassay represented release of genuine bioactive prostacyclin we used a human platelet bioassay(Mitchell *et al*., 2019), analogous to the methodology used in the original identification of prostacyclin (Moncada *et al*., 1976). Addition of lung segments to human platelet-rich plasma resulted in an inhibition of stimulated platelet aggregation; this aggregation was absent when lung from global cyclo-oxygenase-1-deficient mice was used, consistent with an anti-platelet effect of lung-derived prostacyclin (Figure 3I). By contrast, lung from endothelial-selective or endothelial/platelet cyclo-oxygenase-1-deficient mice retained their anti-platelet activity in full agreement with idea that prostacyclin generation in lung occurs independently of the endothelial cyclo-oxygenase/prostacyclin synthase pathway (Figure 3I).

### Endothelial cells from the lung are deficient in prostacyclin synthesis compared to those from large arteries

The lung is amongst the most highly vascularised organs in the body. As such, to understand the presence of a ‘non-vascular’ prostacyclin pathway we had to first determine why lung endothelial cells do not meaningfully contribute to prostacyclin production. Studies of isolated, cultured cells certainly show that lung microvascular endothelial cells are capable of prostacyclin synthesis. However, to our knowledge the relative capacity of lung endothelial cells to synthesise prostacyclin has not been systemically compared, head-to-head, with endothelial cells from large arteries. Further, it is well established that, in culture, there are rapid changes in endothelial expression and activity of cyclooxygenase and prostacyclin synthase; changes that confound extrapolation of prostanoid release patterns in culture to those in the body(Ager *et al*., 1982). Therefore, to address this issue we isolated fresh, matched endothelial cells (CD31^+^, CD45^-^, CD41^-^) from mouse aorta and lung by FACS (Figure 4A). These cells were tested for prostacyclin release immediately after isolation, in the presence of cycloheximide to limit changes in prostanoid synthetic enzymes during their preparation (<4 hours). Endothelial cells from lung, whilst capable of producing prostacyclin, did so at a much reduced level in comparison to aortic endothelial cells (Figure 4B). The suggestion that endothelial cell phenotypes vary according to anatomical location is not new; endothelial cells from the lung microvasculature carry distinct molecular and functional signatures reflecting a more immature and stem cell-like phenotype in comparison to endothelial cells from arteries and veins (Alvarez et al., 2008). Moreover, single cell RNA sequencing studies of endothelial heterogeneity identified prostacyclin synthase in the top 20 transcripts differentiating human lung arterial and capillary endothelial cells(Kalucka et al., 2020). To translate our findings from mice to man we obtained matched, histologically normal pulmonary artery and lung parenchyma from human donors undergoing lung resection for carcinoma and isolated endothelial cells from each using the same approach described for mouse endothelial cells (Figure 4C). As observed for these studies from mice, endothelial cells from human lung parenchyma showed the same pattern of lesser prostacyclin synthesis when compared to endothelial cells isolated from pulmonary arteries of the same individuals (Figure 4D)

**Figure 4.**
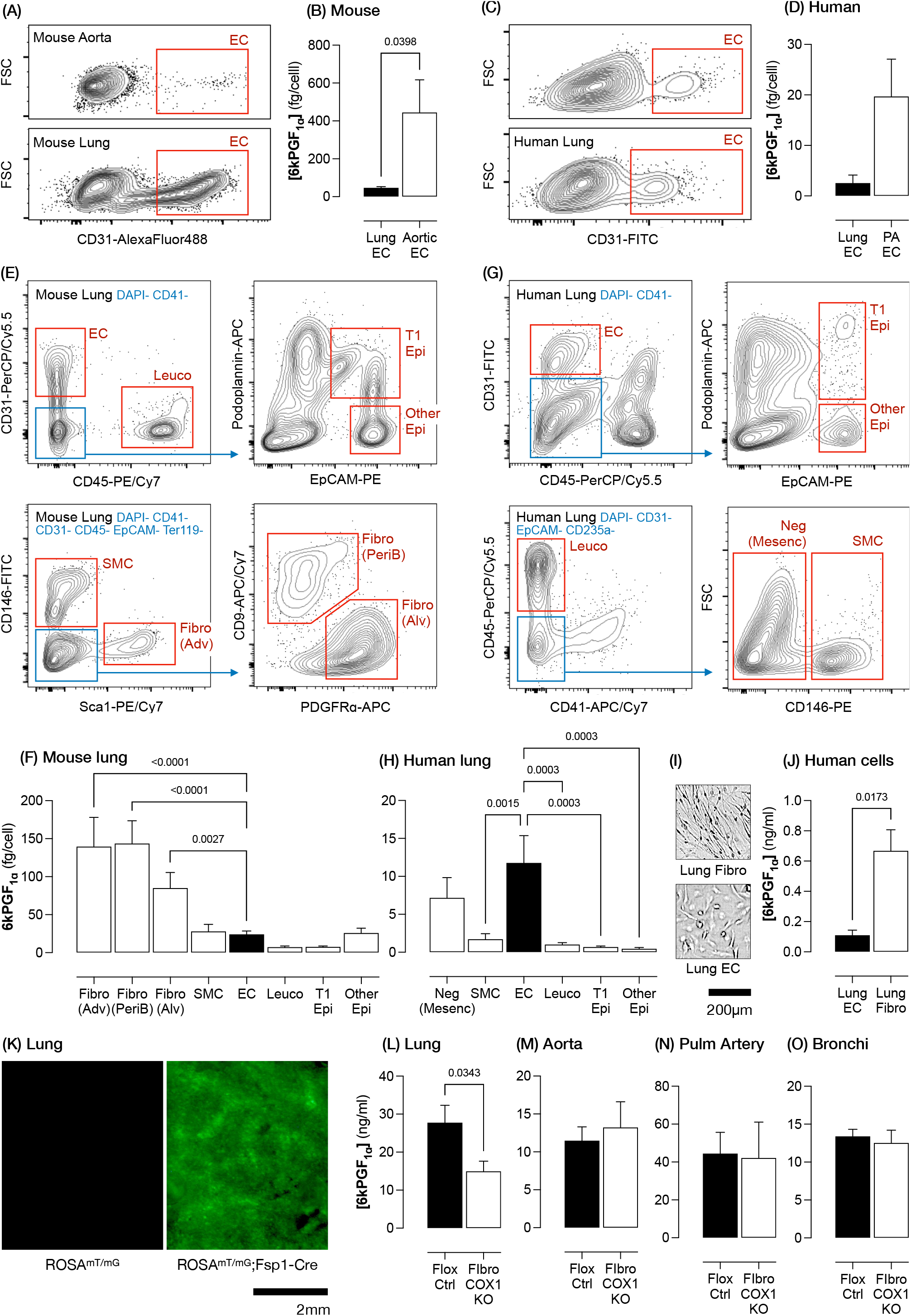
Fibroblasts are major contributors to lung prostacyclin generation. (A) FACS gating strategy (representative plots) and (B) prostacyclin release (measured as 6kPGF_1α_ after arachidonic acid 30μM stimulation) (n=4) of endothelial cells from matched mouse aorta and lung parenchyma. (C) FACS gating strategy (representative plots) and (D) prostacyclin release (n=4 independent isolations from n=2 donors) of endothelial cells from matched human pulmonary artery and lung. In each, endothelial cells were defined as DAPI-, CD41-, CD45-, CD31+ events. (E) FACS gating strategy (representative plots) and (F) prostacyclin release from endothelial cells (EC), leucocytes (Leuco), type 1 (T1 Epi) and other epithelial cells (Other Epi), smooth muscle cells (SMC) and adventitial (Fibro Adv), alveolar (Fibro Alv) and peribronchial fibroblasts (Fibro PeriB) from mouse lung (n=6-12). (G) FACS gating strategy (representative plots) and (H) prostacyclin release from EC, Leuco, T1 Epi, Other Epi, SMC and negatively selected mesenchymal cells (Neg Mesenc) from human lung (n=7-11 donors). (I) Representative brightfield images and (J) prostacyclin release (after arachidonic acid 30μM stimulation) (n=3 donors) from cultured primary human lung microvascular endothelial cells (Lung EC) and human lung fibroblasts (Lung Fibro). (K) EGFP fluorescence (green) in lung segments of ROSA^mT/mG^ mice with/without a Fsp1-Cre transgene (representative of n=3/genotype). Prostacyclin release (after A23187 Ca^2+^ ionophore 30μM stimulation) from (L) lung parenchyma segments (n=12-16), (M) aorta (n=12-16), (N) pulmonary artery (n=6-7) and (O) bronchi (n=6) from fibroblast-specific cyclo-oxygenase-1 knockout (Fibro COX1 KO) and floxed littermate control mice (Flox Ctrl). Plots show 5% density contours. Data are mean ± SEM with p values by repeated measures one-way ANOVA with Holm-Sidak post-test (F,H) or unpaired (B, D, J, L-O) t-test indicated where p<0.05.

### Fibroblasts are the principal contributors to prostacyclin production in the lung

If endothelial cells are not the major source of lung prostacyclin, what cell type(s) account for its production? This question cannot be addressed by immunohistochemical/gene expression approaches, both because of the complex cascade of enzymes and biochemical factors required to support prostacyclin synthesis and because of limitations in the specificity/sensitivity of antibodies to relevant target proteins. To answer this question we again studied cell populations rapidly FACS-isolated from fresh mouse lung tissue. Using endothelial cells (CD31^+^ CD45^-^) as a benchmark, we profiled prostacyclin release from epithelial cells (EpCAM^+^, Podoplanin^-^), type I alveolar epithelial cells (EpCAM^+^, Podoplanin^+^) and leucocytes (CD45^+^) (Figure 4E), isolated by the methodology described by Funino et al(Fujino et al., 2012). From the same mice we also isolated a mesenchymal cell population by exclusion (EpCAM^-^, CD45^-^, CD31^-^, Ter119^-^) and, within this cell population, used an approach defined from single cell RNAseq analysis of the lung (Tsukui et al., 2020) to positively select for smooth muscle cells (CD146^+^), adventitial fibroblasts (CD146^-^ Sca1^+^), alveolar fibroblasts (CD146^-^ Sca1^-^ PDGFRα^+^) and peribronchial fibroblasts (CD146^-^, Sca1^-^, PDGFRα^-^, CD9^+^) (Figure 4E). When stimulated with arachidonic acid to maximally activate cyclo-oxygenase pathways, the leucocytes, epithelial cells and smooth muscle cells released comparable (or numerically less) prostacyclin than endothelial cells (Figure 4F). However, fibroblasts of each of the isolated subtypes exhibited significantly greater prostacyclin production (up to 6-fold greater than endothelial cells; Figure 4F). We replicated these studies with cells isolated from fresh, histologically normal human lung tissue. Endothelial cells (CD31^+^ CD45^-^), pithelial cells (EpCAM^+^, Podoplanin^-^), type I alveolar epithelial cells (EpCAM^+^, Podoplanin^+^) and leucocytes (CD45^+^) were defined and isolated in the same fashion as from mouse lung (Figure 4G). Because suitable cell surface markers to positively select human lung fibroblast populations have not been defined, we studied only a negatively selected ‘mesenchymal cell’ population (EpCAM-, CD45^-^, CD31^-^, CD235a^-^) which was subdivided into smooth muscle cells (CD146^+^) and other cells (CD146^-^) (Figure 4G). Prostacyclin release from leucoyctes, epithelial cells and smooth muscle was low, when compared to endothelial cells (Figure 4H). Prostacyclin levels released from negatively selected mesenchymal cells were similar to those from endothelial cells (Figure 4H). These data are consistent with the suggestion that, in human lung, both fibroblasts and endothelial cells are meaningful contributors to total prostacyclin release. However this negative selection approach may underestimate fibroblast prostacyclin synthesis, as a consequence of possible contamination by other cell types. To explore this question further we performed similar experiments using commercially sourced primary human lung cell cultures. When grown under identical conditions, human lung fibroblasts and microvascular endothelial cells exhibited morphologies characteristic of their respective cell lineage (Figure 4I). After arachidonic acid stimulation both cell types produced prostacyclin; however, the prostaglandin levels released from primary human lung fibroblasts were greater than those from primary lung microvascular endothelial cells (Figure 4J). Although these data are interpreted with some caution, given the changes in prostanoid pathways that occur in vitro, the results support our findings from freshly isolated mouse and human cells that fibroblasts are central contributors to lung prostacyclin production.

Having identified fibroblasts as the major prostacyclin-producing cells in mouse and human lung, we returned to a mouse cell-specific knockout approach to understand the fibroblast contribution to bulk tissue prostacyclin release. Fibroblasts are a heterogeneous cell type; no one marker/promoter identifies the full spectrum of cells defined as fibroblasts. Consequently, the selection of an appropriate model mouse model is difficult. We chose to generate a Cre/loxP fibroblast knockout model driven by the fibroblast-specific protein-1 (Fsp1/S100a4) promoter. Fsp1 is expressed in lung fibroblasts(Iwano et al., 2002). Although, this promoter is also active in monocyte/macrophages, these cells have almost no prostacyclin synthetic capacity (Figures 4F, 4H), suggesting there should be little or no substantial consequences of ‘off-target’ deletions here. To confirm the suitability of this model, we crossed Fsp1-Cre mice with EGFP reporter animals and observed robust recombination within the lung (Figure 4K). When Fsp1-Cre mice were crossed onto a Ptgs1^flox/flox^ background to generate fibroblast-specific cyclo-oxygenase-1 knockout mice (Ptgs1^flox/flox^;Fsp1-Cre), a ~50% reduction occurred in stimulated prostacyclin release from intact segments of lung parenchyma when compared to floxed littermate controls (Figure 4L). Moreover, fibroblast cyclo-oxygenase-1 deletion had no effect on prostacyclin release from the aorta (Figure 4M), lung vasculature (Figure 4N) or airways (Figure 4O). These data provide strong support for our observations from mouse and human cells and support the hypothesis that fibroblasts are responsible for a substantial component of lung tissue prostacyclin production. The residual prostacyclin synthesis observed in these mice may be attributed to other cell types or sub-sets of fibroblasts which do not express the Fsp1-Cre transgene. Fibroblast-specific cyclo-oxygenase-1-deletion also showed a strong trend to reduce prostacyclin release from gut tissue (colon; p=0.07), which, like the lung, exhibited prostacyclin release essentially independent of the endothelial cyclo-oxygenase-1 (Supplementary Table 2). By contrast, no effect of fibroblast-specific cyclo-oxygenase-1 knockout on prostacyclin release was observed from heart, kidney or spleen (Supplementary Table 2).

### Non-vascular prostacyclin can enter the circulation to provide systemic anti-thrombotic tone

With the data above suggesting that fibroblasts and other non-vascular cells are major contributors to tissue prostacyclin production we considered what biological significance of prostacyclin produced outside the vascular wall. Specifically, we explored whether the prostacyclin produced by fibroblasts in tissues acts simply as a local mediator and/or can access the circulation to produce systemic effects. To address this question we first used an ex vivo isolated perfused lung preparation in which released prostacyclin is collected through the organ’s vasculature. In this model, prostacyclin release (detected as its hydrolysis product 6keto-PGF_1α_) was readily detectable in the perfused tissue effluent (Figure 5A) in agreement with previous work(de Deckere *et al*., 1977). These levels were slightly, but significantly, reduced in endothelial/platelet cyclo-oxygenase-1 knockout mice (Figure 5A) consistent with our finding that arterial endothelial cells are effective prostacyclin producers (Figure 4). Notably, perfused lung from fibroblast-specific cyclo-oxygenase-1 also showed a strong reduction in prostacyclin levels in the venous outflow (Figure 5A). In agreement, 6keto-PGF_1α_ levels in plasma were reduced both by endothelial/platelet and by fibroblast cyclo-oxygenase-1 deletion models, when compared to their respective control strains (Figure 5B). Thus, prostacyclin derived from endothelial cells and prostacyclin derived from fibroblasts can both be detected as 6keto-PGF_1α_ in the vascular compartment.

**Figure 5.**
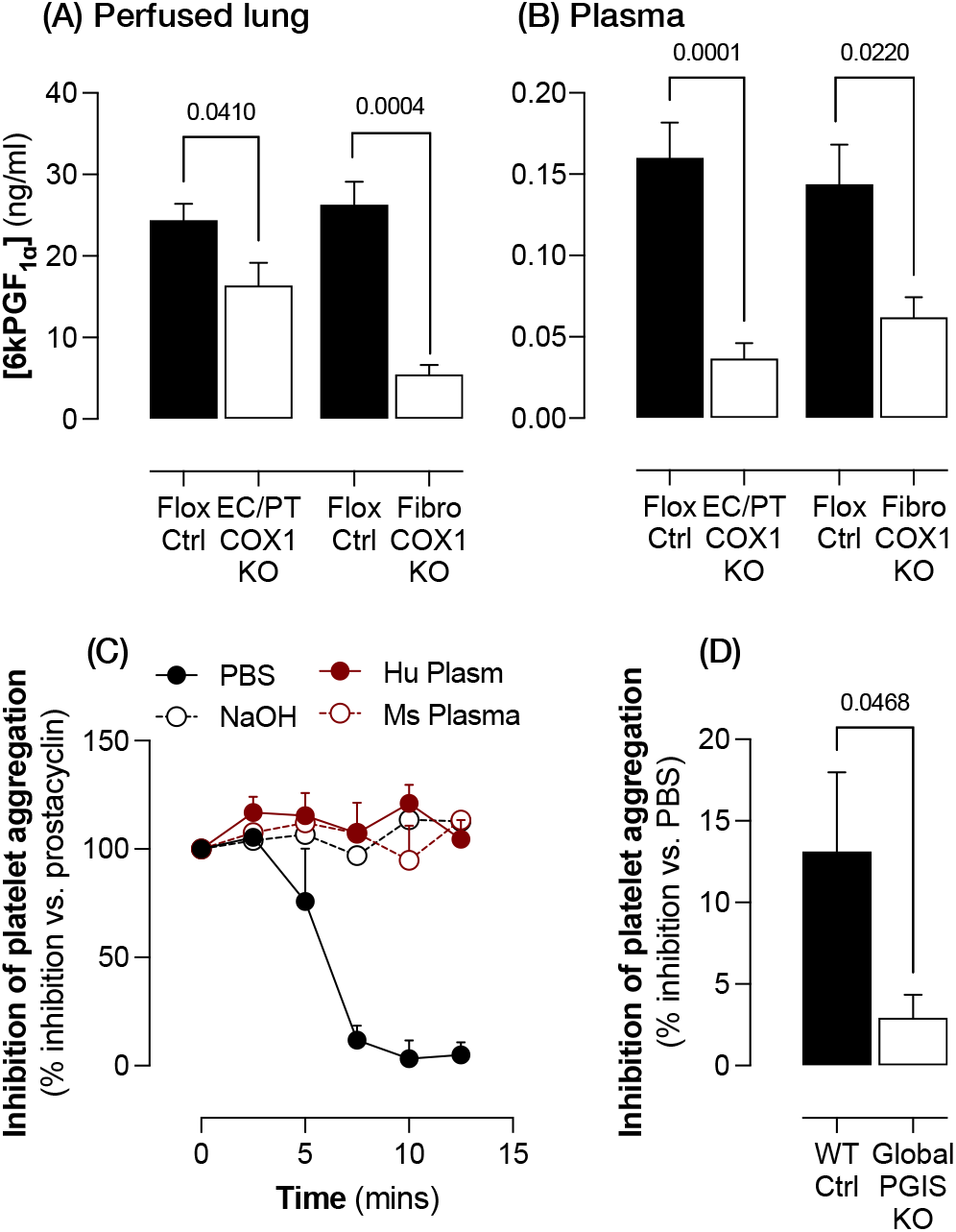
Prostacyclin from fibroblasts and endothelial cells enters the systemic circulation. Prostacyclin levels (measured as 6kPGF_1α_) measured in the outflow from (A) isolated perfused lung (n=4-19) and (B) plasma (n=9-15) from endothelial/platelet cyclo-oxygenase-1 knockout (EC/PT COX1 KO), fibroblast cyclo-oxygenase-1 knockout (Fibro COX1 KO) and floxed control animals (Flox Ctrl). (C) Effect of exogenous prostacyclin (10nM) pre-incubated at 37C for indicated times in PBS, NaOH (0.01M), human (Hu plasma) or mouse plasma (Ms Plasma) (n=3-4) or (D) of plasma from wild-type control (WT Ctrl) and global prostacyclin synthase knockout mouse plasma prepared within 3 mins of collection (n=12) on U46619-stimulated human platelet aggregation. Data are mean ± SEM with p values by unpaired (A,B) or paired (D) t-test indicated where p<0.05.

While it is clear that fibroblast-derived 6keto-PGF_1α_ can be found in the vascular system, the critical question remains: does fibroblast-derived prostacyclin enter the vascular system in an active form to exert an anti-platelet/vascular protective effect or is it present only as an its inactive breakdown product? Endothelial cells interface with the blood so prostacyclin released in vessels will be well-placed to exert local antiplatelet effects. However, for fibroblasts, which do not interface with blood directly, any fibroblast-derived prostacyclin would need to enter the circulation to have a systemic anti-platelet effect *in vivo*. The idea that prostacyclin ‘circulates’, rather than acts simply as a local mediator, is plausible but not universally accepted. The controversy arises because the stability of prostacyclin is dependent upon the matrix in which it is located. Prostacyclin has a half-life of 2-3 mins in physiological buffers, but up to 3 and 15 min in blood or plasma(Dusting et al., 1978), which compares favourably to the < 1min it takes blood to make a complete passage of the circulation. We addressed this directly by comparing the stability of exogenous prostacyclin in PBS (pH7.4), mouse and human plasma and 10mM NaOH, a buffer known to stabilise prostacyclin. The stability of prostacyclin was measured by bioassaying the solutions for their ability to inhibit aggregation of human platelets (Figure 5C). 10nM prostacyclin, a maximal inhibitory concertation, lost all anti-platelet activity when pre-incubated at 37°C in PBS for between 5 and 7.5 mins (Figure 5C). When the preincubation was carried out in human plasma, mouse plasma or 10mM NaOH, however, no loss of anti-platelet activity was observed up to 12.5 mins (Figure 5C). These data are consistent with previous reports demonstrating the presence in blood of ‘prostacyclin stabilising factors,’ such as apolipoprotein A-I, that protect bioactive prostacyclin from inactivation(Yui et al., 1988). If our in vitro data is reflective of in vivo conditions, freshly isolated plasma should contain biologically active prostacyclin. To test this hypothesis, we took the same bioassay approach. To optimise the experiment and take account of any non-prostacyclin effects of plasma on platelet aggregation, we used samples from global prostacyclin synthase deficient mice; these animals exhibit a complete loss of plasma 6keto-PGF_1α_(Kirkby *et al*., 2020). Addition of wild-type mouse plasma, within 3 minutes of collection, to human platelets resulted in a small (~15%), but measurable suppression of platelet aggregation (Figure 5D). This effect was almost abolished when plasma from global prostacyclin synthase knockout mice was used (Figure 5D). These observations are the first to demonstrate that bioactive prostacyclin is present and circulates in blood and corroborate the idea that non-endothelial derived prostacyclin could impact on thrombotic tone.

If fibroblasts and other non-vascular cells produce prostacyclin that can enter the vascular compartment and have the potential to circulate in a bioactive form, we considered whether these previously unconsidered prostacyclin depots can also contribute actively to cardiovascular protection, in collaboration with endothelium-derived prostacyclin. Prostacyclin is best understood as an anti-platelet/anti-thrombotic factor; consequently, we have previously used an in vivo FeCl_3_ carotid artery injury model to demonstrate a pro-thrombotic phenotype in endothelium-specific cyclo-oxygenase-1 knockout mice(Mitchell *et al*., 2019). Here, we used the same model to show that fibroblast-specific cyclo-oxygenase-1 knockout mice also exhibited a modest but significant pro-thrombotic phenotype – showing reduced mean occlusion time compared to floxed littermate control mice (Figure 6A,B). This result was not associated with any change in local carotid artery prostacyclin generation (Flox Ctrl: 33.1±9.5ng/ml; Fibro COX1 KO: 34.5±23.3ng/ml; n=6; p>0.05 by unpaired t-test), which we have previously shown to be predominately generated by endothelial cells(Mitchell *et al*., 2019). Thus, platelet inhibitory prostanoids (presumably prostacyclin) derived from fibroblasts in lung and/or elsewhere also provide systemic anti-thrombotic protection.

**Figure 6.**
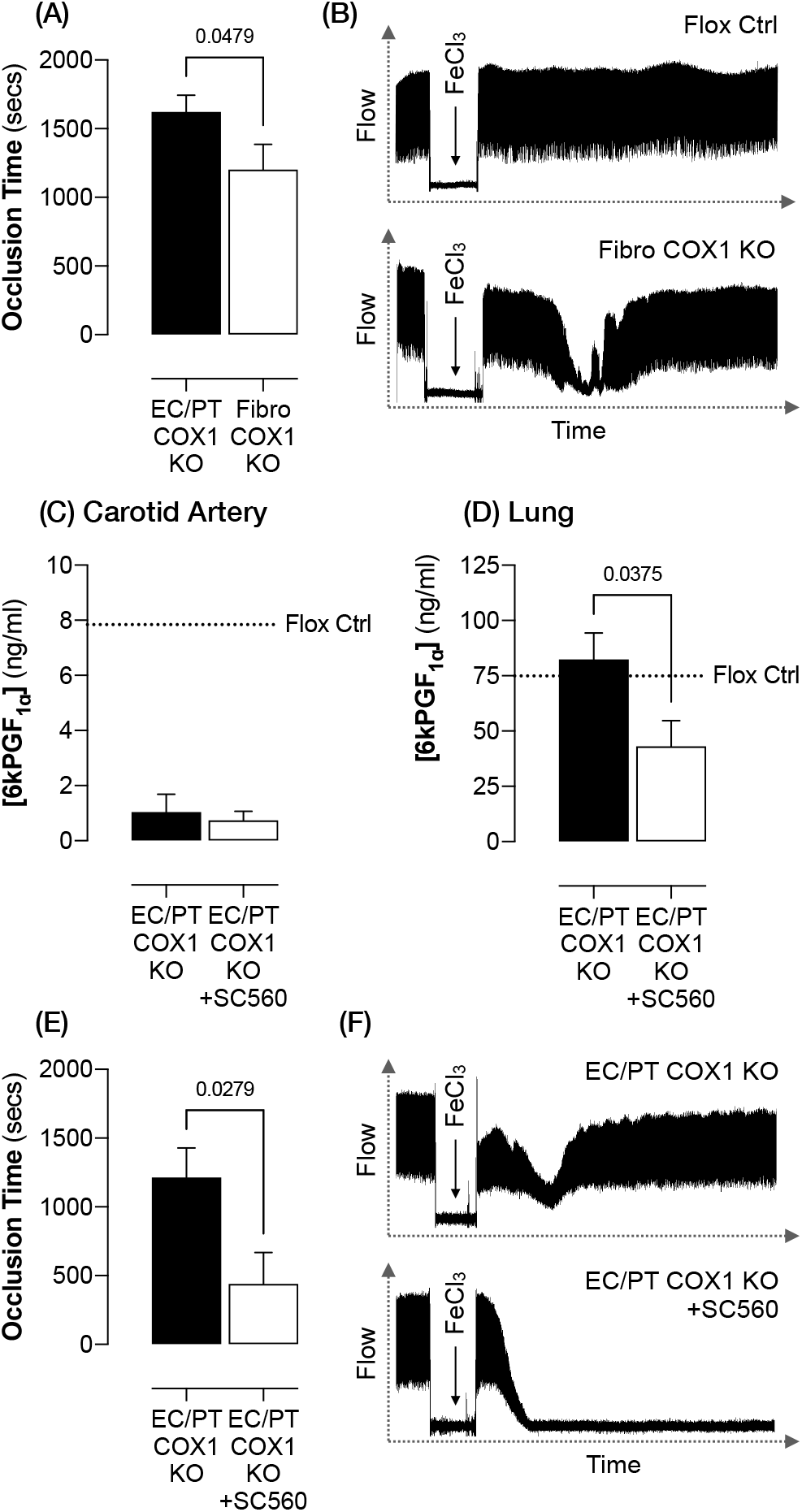
Fibroblasts and other non-vascular sites of cyclo-oxygenase-1 expression contribute to systemic anti-thrombotic protection. (A) Thrombotic occlusion time (n=14-15) and (B) representative blood flow traces after carotid artery FeCl_3_ injury in vivo in fibroblast cyclo-oxygenase-1 knockout mice (Fibro COX1 KO) and floxed littermate controls (Flox Ctrl). Prostacyclin release (measured as 6kPGF_1α_) from (C) carotid artery (n=7) and (D) lung parenchyma (D; n=7-9) ex vivo from endothelial/platelet cyclo-oxygenase-1 knockout mice (EC/PT COX1 KO) treated with the cyclo-oxygenase-1 inhibitor, SC-560 (10mg/kg; iv, 15 mins) or vehicle (5% DMSO). Release level from Flox Ctrl tissue is marked on each graph as a dashed line. (E) Thrombotic occlusion time (n=7-13) and (F) representative blood flow traces after carotid artery FeCl3 injury in vivo in EC/PT COX1 KO with treated with SC-560 or vehicle. Data are mean ± SEM with p values by Mann-Whitney U-test (A,E) or unpaired t-test (C,D) indicated where p<0.05.

Fibroblast-specific cyclo-oxygenase-1 knockout mice exhibit only a partial loss of tissue prostacyclin production, suggesting there may be other sources that contribute to the total anti-thrombotic contribution of all non-vascular prostacyclin sources of in the body. To approach this we determined the effect of pharmacologically removing all residual sources of prostacyclin in mice already lacking vascular and platelet prostanoids. To this end, we treated endothelial/platelet cyclo-oxygenase-1 knockout mice with the selective cyclo-oxygenase-1 inhibitor, SC-560, to determine the effect of removal of cyclo-oxygenase-1-derived prostanoids from all non-vascular cells. Because these mice already lack cyclo-oxygenase-1 in platelets and endothelial cells, no effect of SC-560 on carotid artery prostacyclin levels (Figure 6C) or platelet thromboxane levels was noted (Supplementary Figure 1) but prostacyclin release by lung tissue was reduced ~50% (Figure 6D). This prostacyclin reduction was associated with a marked increase in thrombosis after carotid artery FeCl_3_ injury (Figure 6E,F). This effect could not be attributed to an off-target SC-560 effect, because this treatment had no effect on thrombosis in global cyclo-oxygenase-1-deficient mice (Supplementary Figure 2).

## Discussion

The identification of prostacyclin as an endogenous mediator with profound anti-platelet activity was rapidly followed by the idea that manipulating prostacyclin could offer the means to prevent and treat cardiovascular disease(Mitchell and Kirkby, 2019; Moncada *et al*., 1976). In the 45 years since these first observations, this therapeutic potential has remained largely untapped; to date prostacyclin therapy has found important, but only relatively niche, use as a treatment for pulmonary hypertension. This failure to realise the potential of targeting prostacyclin biology likely reflects the subsequent discovery of a great deal of complexity underlying its biology, including multiple synthetic pathways(McAdam *et al*., 1999), competing local and systemic actions(Kirkby *et al*., 2013b; Mitchell *et al*., 2019) and promiscuous receptor pharmacology(Liu et al., 2012a; Mitchell *et al*., 2021). Both difficulties of measuring prostacyclin synthesis in vivo(Mitchell *et al*., 2018) and side effects associated with systemic prostacyclin delivery also complicate exploiting prostacyclin biology for therapeutic purposes. Since its initial discovery, the idea that vascular endothelium is the major source of prostacyclin in the body has remained largely intact. Our current findings however suggest this too must be re-evaluated, the data presented herein demonstrate a role for fibroblasts and other non-vascular cells in both prostacyclin synthesis and consequent systemic anti-thrombotic protection. Whilst prostacyclin production by non-vascular cells is not in itself a novel concept, to our knowledge this report constitutes the first demonstration that these other depots of prostacyclin production make a meaningful and functionally significant contribution to prostacyclin’s systemic bioactivity. Thus, we must now also consider the biological role and therapeutic potential of non-vascular prostacyclin produced in the lung and elsewhere.

Firstly, it is clear that pulmonary vascular disease is associated with prostacyclin deficiency(Christman et al., 1992) and cardiovascular risk. These new findings raise the possibility that diseases of the lung parenchyma may also be associated with a loss of cardioprotective prostacyclin. For example, both chronic obstructive pulmonary disorder and interstitial lung disease feature dysfunction, damage, and phenotypic alterations to a range of lung parenchymal cells, including fibroblasts. An impairment of the ability of these non-vascular cells to produce prostacyclin could not only contribute to disease pathogenesis but also explain the marked increased in atherothrombotic risk observed in both conditions(Clarson et al., 2020; Feary et al., 2010). Indeed, lung tissue from chronic obstructive pulmonary disorder patients exhibits reduced prostacyclin synthase expression(Nana-Sinkam et al., 2007). Similarly, fibroblasts isolated from pulmonary fibrosis patients show reduced prostacyclin and increased thromboxane formation relative to fibroblasts from healthy controls(Wilborn et al., 1995). These data suggest that a prostacyclin-based therapy may help to mitigate the excess cardiovascular risk and/or treat disease progression in these specific patient groups. In support of this suggestion, inhaled trepostinil has therapeutic benefits in in the treatment of idiopathic pulmonary fibrosis(Harari and Wells, 2021). Similar hypotheses could also be formulated for non-vascular diseases of other organ systems associated with increased risk of thrombotic events.

Secondly, our finding that lung microvascular endothelial cells in vivo are relatively deficient in prostacyclin production may also suggest an opportunity to boost its endogenous generation to treat both lung and thrombotic disease. Over-expression of prostacyclin synthase in the lung by various means is protective in animal disease models of pulmonary hypertension(Geraci et al., 1998). Whilst this non-specific delivery is clearly effective, within the lung, endothelial cells are naturally positioned to respond to local vascular conditions and to deliver hormones. In light of our findings we suggest that, to maximise efficacy and reduce the systemic side effects which plague prostacyclin therapies, prostacyclin synthase delivery approaches would best be targeted to the pulmonary endothelium. Recently developed simple polymer-based transfection reagents that selectively deliver mRNA cargoes to lung endothelium after systemic administration(Kaczmarek et al., 2018) could potentially provide a practical route to this approach.

Thirdly, we have focussed here on the idea that prostacyclin derived from fibroblasts or from other non-vascular sources acts as a systemic biological mediator. However, we must also consider the possibility that fibroblast-derived prostacyclin may have important autocrine or paracrine roles. Prostacyclin has well-defined effects on lung cell function; effects that include bronchodilation, immunomodulation, and inhibition both of fibrosis and proliferation. All these responses may be associated, partly or wholly, with prostacyclin derived from non-vascular cell sources. The alternative cellular sites of prostacyclin production and how prostaglandin production at these sites is altered in lung disease should now be considered, to determine whether therapeutic targeting of these sites and/or modulation of their prostaglandin production could be exploited to improve lung function and/or protect against development of lung disease.

Whilst our observations are limited to preclinical studies, the idea that there are meaningful non-vascular prostacyclin sources presents a new concept in our understanding of prostacyclin biology. This is not suggested to reduce emphasis on the importance of endothelial cyclo-oxygenase-1(Mitchell *et al*., 2019) and cyclo-oxygenase-2-derived prostacyclin(Mitchell *et al*., 2019) and the consequences of these powerful anti-thrombotic pathways – the evidence of their importance is clear – but we must now also consider a parallel cardioprotective and/or local disease modifying pathway associated with cyclo-oxygenase-1 and prostacyclin in non-vascular cells and tissues. In principle, each of these pathways generates the same prostanoid products; but there appears to be a lack of redundancy such that each of these alternative sources of prostacyclin carries unique biological functions. Consequently, loss of any individual prostacyclin production source/pathway may have profound consequences for cardiovascular health. The roles of these alternative sources of prostacyclin production, individually and together, should now be evaluated in context, through health and disease and across organ systems. These results suggest new therapeutic opportunities, within these new prostacyclin pathways, for the treatment of a range of human diseases, including cardiovascular disease as a co-morbidly of respiratory conditions.

## Methods

### Animals

Studies were performed on 8-12 week old male and female mice housed in individually ventilated cages with free access to standard chow and water and 12h day/night cycle. All procedures were conducted in accordance with the UK Animals (Scientific Procedures) Act (1986) Amendment (2013) and the Guide for the Care and Use of Laboratory Animals published by the US National Institutes of Health (NIH Publication No. 85-23, revised 1996) and after local approval from the Imperial College Animal Welfare Ethical Review Board (UK Home Office License Project Licenses 70/7013 and PP1576048), the Institutional Animal Research and Use Committees of Shantou University (approval SUMC2019-336) or Fudan University. Global cyclo-oxygenase-1 knockout mice (*Ptgs1*^-/-^)(Langenbach et al., 1995) and global prostacyclin synthase knockout mice (*Ptgis*^-/-^)(Kirkby et al., 2020) were generated as previously described and compared to age and sex matched wild-type C57Bl/6 controls (Charles River, UK or Vital-River, China). Endothelial-specific cyclo-oxygenase-1 knockout mice (*Ptgs1^flox/flox^; VEC-iCre*)(Mitchell *et al*., 2019), endothelial/platelet-specific cyclo-oxygenase-1 knockout mice (*Ptgs1^flox/flox^; Tie2-Cre*)(Mitchell *et al*., 2019), endothelial/platelet-specific cyclo-oxygenase-2 knockout mice (*Ptgs2^flox/flox^; Tie2-Cre*)(Mitchell *et al*., 2019) and endothelial/platelet-specific prostacyclin synthase (*Ptgis^flox/flox^; Tie2-Cre*)(Cao et al., 2019) were generated as we have previously described. Because the *VEC-iCre* used to generate endothelial cyclo-oxygenase-1 knockout mice requires activation by tamoxifen, at 4-6 weeks of age, all mice from this line were treated with tamoxifen (50mg/kg, ip, once daily for 5 days; Sigma, UK) and allowed to recover to 2 weeks before further use. Fibroblast-specific cyclo-oxygenase-1 knockout mice (*Ptgs1^flox/flox^; Fsp1-Cre*) were generated by crossing floxed *Ptgs1* mice (JAX strain: 30884)(Mitchell *et al*., 2019) with transgenic *Fsp1/S1004A4-Cre* mice (JAX strain: 12641)(Tsutsumi et al., 2009) and were maintained on a mixed C57Bl/6-BALB/c background. EGFP/Cre activity reporter strains were generated by crossing ROSA^mT/mG^ mice (JAX strain: 7676)(Muzumdar et al., 2007) with transgenic Tie2-Cre (JAX strain: 8863)(Kisanuki et al., 2001) or Fsp1/S100A4-Cre mice and were studied as heterozygous animals. For all cell-specific knockout strains, animals were genotyped by genomic PCR to identify Cre-positive knockout animals from the Cre-negative littermates which were used as experimental controls for each strain. Unless otherwise indicated, animals were euthanised by CO_2_ narcosis. Animal models are available on reasonable request to the authors.

### Human samples

All studies using human material were conducted in accordance with the Declaration of Helsinki and samples were donated from volunteers and patients who gave explicit informed consent. Blood was collected into sodium citrate (0.32% final; BD Biosciences, Germany) from healthy male and female volunteers aged 18-40 after ethical approval by the West London & GTAC Research Ethics Committee (approval 15/LO/223), the St Thomas’ Hospital Research Ethics Committee (approval 07/Q0702/24) or the Institutional Ethic Committee of Shantou University (approval SUMC-2019-47). Lung parenchyma and pulmonary artery was collected from patients undergoing surgical lung resection for treatment adenocarcinoma, small cell carcinoma or squamous carcinoma. Histologically normal areas of the resected tissue was identified for study by a clinical pathologist. 3 male and 7 female patients with an average age 73.7 (range 69-87) donated lung parenchyma samples. Of these, 1 male and 1 female patient with an average age of 74.5 (range 71-78) also donated pulmonary artery. Lung tissue was provided through the Royal Brompton & Harefield NHS Trust Biomedical Research Unit Advanced Lung Disease Biobank after ethical approval by the South Central – Hampshire B Research Ethics Committee (approval 15/SC/0101) and local project review by the Biomedical Research Unit Heads of Consortia (approval JM04).

### Tissue and plasma prostanoid levels

#### Tissue/vessel segments

Tissue segments (approximately 10mm^3^) or vascular rings (approximately 2mm long) were cleaned of adherent material, transferred to 96-well microplate wells containing A23187 (Ca^2+^ ionophore; Sigma, UK; 30μM) or arachidonic acid (Sigma, UK; 30μM) in DMEM media (Sigma, UK). After 30 mins incubation at 37°C supernatant was collected. Where indicated, before tissue collection the lungs were flushed of blood by perfusion of the pulmonary vasculature with PBS via the right ventricle with effective perfusion confirmed by blanching of the lung tissue.

#### Blood

Blood was collected from the inferior vena cava into heparin (10U/ml final; Leo Laboratories, UK) and stimulated with A23187 (30μM) for 30 mins, before centrifugation (8000g, 2 mins) and separation of conditioned plasma.

#### Homogenates

Segments of lung parenchyma were removed and snap frozen. Lung segments were suspended in 10X volume of ice cold PBS containing cOmplete Mini protease inhibitor cocktail (Roche, Switzerland), 2mM EDTA (Sigma, UK) and an excess of the non-selective cyclo-oxygenase inhibitor, diclofenac (1mM; Sigma, UK). Samples were immediately homogenised using a Precellys24 instrument (Bertin Instruments, France) and the supernatant collected after centrifugation (8000g; 2 mins).

#### Plasma

Blood was collected from the inferior vena cava into heparin (10U/ml final) immediately postmortem. Plasma was separated by centrifugation (8000g, 2 mins) and stored.

#### Perfusates

Immediately post-mortem, the thoracic cavity was opened and the pulmonary vasculature flushed of blood with PBS via the right ventricle. The right atrium of the heart was then cannulated with PE10 tubing, secured with 5-0 silk and the lung and heart were removed intact. The pulmonary vasculature was perfused via the right atrium with DMEM media at 37°C for 20 mins using a peristaltic pump at 50μl/min and the venous outflow collected from the left atrium.

Levels of the prostacyclin breakdown product 6-keto-PGF_1α_ (Cayman Chemical, USA) and/or PGE2 (Biotechne, UK) were measured in supernatants/plasma by commercial immunoassay. In some cases tissue/vessel segments were weighed and prostanoid levels expressed relative to tissue mass.

### qPCR

Tissue segments were snap frozen, then homogenised in ice cold RLT buffer (Qiagen, UK) containing b-mercaptoethanol (1% v/v; Sigma, UK) using a Precellys24 instrument. RNA was extracted using RNeasy mini-prep kits (Qiagen, UK) and gene expression levels measured using a 1-step RT-qPCR master mix (Promega, UK) and a 7500 Fast qPCR instrument (Applied Biosystems, UK) using TaqMan probes (Qiagen, UK) recognising Ptgis (probe ID: Mm00447271_m1), Ptgs1 (probe ID: Mm00477214_m1), Ptgs2 (probe ID: Mm00478374_m1) or the housekeeping genes 18S (probe ID: Mm03928990_g1) and GAPDH (probe ID: Mm99999915_g1). Data were analysed by the comparative Ct method, with relative expression levels normalised to those to 18S and GAPDH and experimental control groups.

### Platelet prostacyclin bioassay

A human platelet bioassay was used to measure bioactive prostacyclin levels released from segments of mouse lung parenchyma (approximately 10mm^3^) as we have described previously(Mitchell *et al*., 2019). Briefly, platelet-rich plasma (PRP) and platelet-poor plasma (PPP) were separated from human blood by centrifugation (PRP: 230g, 15 mins; PPP: 8000g, 2 mins). PRP was pre-incubated with aspirin (30μM; 30 mins prior; Sigma, UK) and DEA/NONOate (10μM; 1 min prior; Sigma, UK) to sensitise platelets to prostacyclin. Lung segments were added to individual wells of 96 well microtitre plates containing PRP and pre-incubated for 1 min, before stimulation of platelets and tissues with A23187 (30μM) and vigorous mixing (1200RPM; BioshakeIQ, Q Instruments, Germay). After 5 mins, the tissue segments were removed and the absorbance of each well at 595nm measured by spectrophotometer and the amount of platelet aggregation calculated by reference to the absorbance of unstimulated PRP (0% aggregation) or PPP (100% aggregation). Bioassay of mouse plasma and of prostacyclin solutions incubated in different buffers were conducted accordingly to broadly the same method. For plasma, mouse blood was collected from the inferior vena cava into heparin (10U/ml final) and plasma quickly separated by centrifugation (10000g, 1 min) before addition to the bioassay in a volume ratio of 1:4 – the entire procedure was completed within 3 mins of blood collection. For prostacyclin stability experiments, prostacyclin (100nM; Biotechne, UK) was incubated at 37°C for 2.5-12.5 mins in wild-type mouse plasma, human plasma, PBS or NaOH (0.01M; Sigma, UK) before addition to the bioassay. In these both experiments, human PRP was not pre-treated with aspirin or DEA/NONOate and U46619 (1μM; Cayman Chemical, USA) was used as the platelet activator.

### Cell isolation

Mouse lung, mouse aorta, human lung parenchyma and human pulmonary artery were finely minced with scissors in an enzyme cocktail of collagenase I (5mg/ml; Sigma, UK), DNase I (125U/ml; Sigma; UK) and elastase (100μg/ml; Sigma, UK) in PBS containing CaCl2 (2mM; Sigma, UK) and incubated at 37°C with regular mixing until fully digested. Cycloheximide (3μM; Sigma, UK) was added to this an all solutions to prevent artefactual changes in prostanoid pathways during cell isolation. Erythrocytes were lysed using ammonium-chloride-potassium lysis buffer (Life Technologies, UK) and cells treated with FcR blocking antibodies (Biolegend, UK). Cell suspensions were then labelled and sorted according the specific protocol as below. All antibodies were purchased from Biolegend, UK. *Mouse lung vs aorta endothelial cells:* Cells were stained with anti-CD45-PE/Cy7, anti-CD31-AlexaFluor488 and anti-CD41-APC/Cy7 and sorted using a FACSAria III instrument (BD Biosciences, Germany).

#### Mouse lung cell panel

Cell suspensions were divided in half and labelled with one of each of the following antibody cocktails before sorting using a FACSMelody instrument (BD Biosciences, Germany). Antibody mix 1 (endothelial/epithelial/leucocyte): anti-EpCAM-PE, anti-CD41-APC/Cy7, anti-CD45-PE/Cy7, anti-CD31-PerCP/Cy5.5 and anti-Podoplanin-APC. Antibody mix 2 (mesenchymal): anti-EpCAM-PE, anti-CD41-PE, anti-CD45-PE, anti-Ter119-PE, anti-CD31-PerCP/Cy5.5, anti-Sca1-PE/Cy7, anti-PDGFRα-APC and anti-CD9-APC/Fire750.

#### Human lung cell panel

Cells from 2 human lung donors were stained with anti-CD41-APC/Fire750, anti-CD45-AlexaFluor488, anti-CD31-BrilliantViolet605, anti-EpCAM-PE and anti-Podoplanin-APC and sorted using a FACSAria III instrument (BD Biosciences, Germany). From these donors ‘Neg (mesenc)’ or ‘SMC’ fractions were not collected. Cells from the remaining human lung donors were divided in half and labelled with one of each of the following antibody cocktails before sorting using a FACSMelody instrument (BD Biosciences, Germany). Antibody mix 1 (endothelial/epithelial): anti-CD41-APC/Cy7, anti-CD45-PerCP/Cy5.5, anti-CD31-FITC, anti-EpCAM-PE and anti-Podoplanin-APC. Antibody mix 2 (mesenchymal/leucocyte): anti-EpCAM-FITC, anti-CD31-FITC, anti-CD235a-FITC, anti-CD45-PerCP/Cy5.5, anti-CD41-APC/Cy7 and anti-CD146-PE.

#### Human pulmonary artery endothelial cells

Cells were stained with anti-CD41-APC/Cy7, anti-CD45-PerCP/Cy5.5 and anti-CD31-FITC and sorted using a FACSMelody instrument (BD Biosciences, Germany).

In each case, single live cells were identified from debris and doublets on the basis of scatter properties and negative DAPI staining and any cells bound to platelets excluded on the basis of CD41 staining. Populations of interest were identified using the gating strategies shown Figures 4 and 5 and approximately 10,000 cells were sorted using 2-way or 4-way purity sort mode using a 100μm nozzle. After separation, cell populations were re-suspended in DMEM media containing arachidonic acid (30μM; Sigma, UK) and incubated at 37°C for 30 mins. The release reaction was stopped by addition of diclofenac (10μM; Sigma, UK) and levels of the stable prostacyclin breakdown product, 6keto-PGF_1α_ measured in the supernatant by immunoassay and expressed relative to cell count.

### Cell culture

Human primary lung microvascular endothelial cells and human primary lung fibroblasts (Promocell, Germany) from 3 individual donors were cultured according to suppliers instructions in full endothelial growth factor-2 media (Promocell, Germany) supplemented with 10% fetal calf serum (Biosera, UK) and penicillin/streptomycin (Sigma, UK). At passage 4-8, cells were plated in 96-well culture plates at a density of 10,000 cells/well in the same media and allowed to settle overnight. The following day, media was replaced and cells stimulated with arachidonic acid (30μM) for 30 mins at 37°C before collection of media for measurement of the stable prostacyclin breakdown product, 6keto-PGF_1α_ by immunoassay. Cells were fixed and counted to confirm the density remained the same between types at the point of stimulation.

### Thrombosis

Thrombosis in vivo was measured using the mouse carotid artery ferric chloride injury model as we have previously described(Mitchell *et al*., 2019; Mitchell et al., 2021). Briefly, under isoflurane anaesthesia the left carotid artery was exposed and separated from the attached nerve and vein. Ferric chloride (4-6%; Sigma, UK) was applied to the adventitial surface of the vessel for 3 mins, then the vessel irrigated and a Doppler peri-vascular flow probe (Transonic Systems, UK) was secured around the artery. Blood flow was recorded for 30 mins and time to occlusion recorded as the time taken from injury to the first point at which blood flow dropped <10% of baseline. If no occlusion occurred, occlusion time was recorded as 30 mins. In some experiments, after anaesthesia, the cyclo-oxygenase-1 inhibitor, SC-560 (10mg/kg; Cayman Chemical, USA) or vehicle (5% DMSO) were administrated intravenously (tail vein) 15 mins prior to arterial injury.

### Statistics & data analysis

Data are presented as mean ± standard error for n individual animals/donors. Where replicate measurements were made from the same individual data were averaged and included as a single data point. Wherever reasonably possible measurements and analysis were performed by an operator blinded to the experimental groups. Statistical analysis was performed using Prism 8 software (GraphPad Software, USA) with tests used indicated in individual figure legends. Data were assumed to be sampled from a normal distribution and therefore parametric tests were used except for thrombosis data in Figure 7 where capping of occlusion time defines a skewed distribution and in this case non-parametric statistics were applied. Differences were considered significant where p<0.05. All data are available in the main text or the supplementary material.

## Acknowledgements

The authors acknowledge Dr Jess Rowley, Dr Larissa Zárate García, Dr Jane Srivastava and the Imperial College South Kensington Flow Cytometry Facility for their assistance in FACS, Mr Harshil Bhayani, Ms Morag Hamilton and the Royal Brompton & Harefield NHS Trust Biomedical Research Unit Advanced Lung Disease Biobank for their assistance in obtaining human lung tissue, Prof Timothy Warner for his provision of global cyclo-oxygenase-1-deficient mice, Mr Gang Yu for his assistance in the provision of global prostacyclin synthase-deficient mice and Dr Changchun Ma and staff at the Cancer Hospital of Shantou University Medical College for their assistance in collecting human blood samples.

## Author Contributions

Investigation & analysis (JAM, MEL-P, FS, PCA, BA-S, YE, BL, NSK), Methodology & models (JAM, BA-S, BL, YZ, C-MH, HRH, NSK), Supervision & Funding: (JAM, BL, YZ, HRH, NSK), Writing – original draft (JAM, NSK), Writing – review & editing (All authors).

