## Supplemental Material for "Tissue fibroblasts are a critical source of prostacyclin and anti-thrombotic protection"

| **[6kPGF$_{1α}$]** (ng/ml) | **Heart** | **Lung** | **Kidney** | **Colon** | **Spleen** |
|---|---|---|---|---|---|
| **Flox Ctrl** | 6.0 ± 2.1 | 53.5 ± 22.0 | 10.0 ± 1.8 | 58.3 ± 19.3 | 14.7 ± 5.3 |
| **EC/PT COX1 KO** | 1.6 ± 0.6 | 45.8 ± 20.2 | 15.7 ± 8.4 | 72.8 ± 23.6 | 9.9 ± 3.2 |
| **Flox Ctrl** | 2.6 ± 0.5 | 30.1 ± 4.6 | 15.3 ± 1.6 | 51.9 ± 7.9 | 28.7 ± 6.6 |
| **EC/PT COX2 KO** | 2.4 ± 0.4 | 39.1 ± 10.0 | 16.3 ± 1.8 | 45.4 ± 7.4 | 21.7 ± 6.1 |
| **Flox Ctrl** | 3.3 ± 1.7 | 13.6 ± 2.5 | 3.3 ± 0.6 | 21.3 ± 3.1 | 3.9 ± 0.5 |
| **EC/PT PGIS KO** | 1.2 ± 0.2 | 17.3 ± 1.1 | 3.2 ± 0.4 | 26.5 ± 4.8 | 4.1 ± 0.5 |

**Supplementary Table 1 Effect of endothelial/platelet cyclo-oxygenase-1, cyclo-oxygenase-2 and prostacyclin synthase deletion on tissue prostacyclin release** Prostacyclin release (measured as 6kPGF$_{1α}$ after A23187 $Ca^{2+}$ ionophore 30μM stimulation) from isolated aorta, heart, lung, renal medulla, renal cortex, colon and spleen from endothelial/platelet cyclo-oxygenase-1 knockout (EC/PT COX1 KO), endothelial/platelet cyclo-oxygenase-2 knockout (EC/PT COX2 KO) and endothelial/platelet prostacyclin synthase knockout mice (EC/PT PGIS KO) each compared to their own respective floxed littermate control mice (Flox Ctrl). n=4-13. Data are mean ± SEM. All p>0.05 by unpaired t-test.

| [6kPGF$_{1\alpha}$] (ng/ml) | Heart | Lung | Kidney | Colon | Spleen |
|---|---|---|---|---|---|
| **Flox Ctrl** | 3.0 ± 0.6 | 27.8 ± 4.5 | 4.6 ± 0.6 | 18.3 ± 2.4 | 8.8 ± 2.0 |
| **Fibro COX1 KO** | 2.9 ± 0.6 | 14.9 ± 2.6 * | 5.4 ± 1.2 | 12.3 ± 2.0 | 6.7 ± 2.1 |

**Supplementary Table 2 Effect of fibroblast cyclo-oxygenase-1 deletion on tissue prostacyclin release** Prostacyclin release (measured as 6kPGF$_{1\alpha}$ after A23187 $Ca^{2+}$ ionophore 30μM stimulation) from isolated heart, lung, kidney, colon and spleen from fibroblast cyclo-oxygenase-1 knockout (Fibro COX1 KO) and floxed littermate control animals (Flox Ctrl). n=9-16. Data are mean ± SEM. *, $p<0.05$ by unpaired t-test.

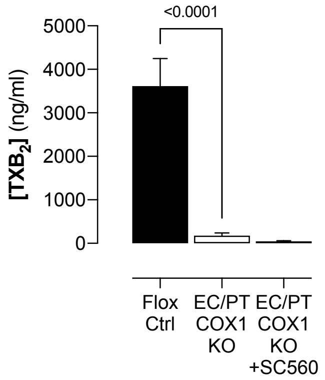


**Supplementary Figure 1 Effect of SC-560 on platelet thromboxane levels in endothelial/platelet cyclo-oxygenase-1 knockout mice** Thromboxane release (measured as thromboxane $B_2$; $TXB_2$; after A23187 $Ca^{2+}$ ionophore 30µM stimulation) from whole blood from endothelial/platelet cyclo-oxygenase-1 knockout mice (EC/PT COX1 KO) treated with the cyclo-oxygenase-1 inhibitor, SC-560 (10mg/kg; iv, 15 mins) or vehicle (5% DMSO) or vehicle-treated floxed littermate control animals (Flox Ctrl). n=5. Data are mean ± SEM with p values by one-way ANOVA with Holm-Sidak post-test indicated where $p<0.05$.

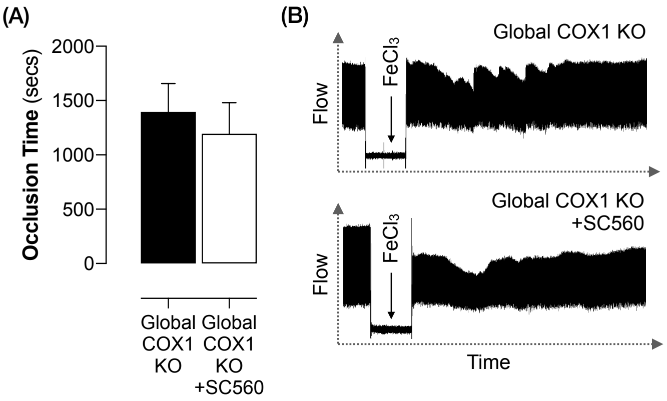


**Supplementary Figure 2 Effect of SC-560 on thrombosis in global cyclo-oxygenase-1 knockout mice** Thrombotic occlusion time (A; n=7) and representative blood flow traces (B) after carotid artery $FeCl_3$ injury in vivo in global cyclo-oxygenase-1 knockout mice (Global COX1 KO) treated with SC-560 (10mg/kg; iv, 15 mins) or vehicle (5% DMSO). Data are mean ± SEM with p values by Mann-Whitney U-test indicated where $p<0.05$.
